# High-Throughput CETSA Identifies Small Molecule Modulators of ILT3 (LILRB4) with Functional Activity in Human iPSC-Derived Microglia for Alzheimers Disease

**DOI:** 10.64898/2026.05.19.726383

**Authors:** Somaya A. Abdel-Rahman, Moustafa Gabr

**Affiliations:** Department of Radiology, Molecular Imaging Innovations Institute (MI3), Weill Cornell Medicine, New York, NY 10065, USA

**Keywords:** CETSA, Alzheimer’s disease, ILT3, Microglia, LILRB4

## Abstract

Immune inhibitory signaling in microglia contributes to impaired amyloid clearance and neuroinflammation in Alzheimers disease (AD), yet small molecule modulators targeting these pathways remain largely unexplored. Here, we report the development of a high-throughput cellular thermal shift assay (HT-CETSA) platform for identification of small molecule binders targeting the inhibitory immune receptor ILT3 (LILRB4). Screening of ∼40,000 compounds yielded multiple validated hits, including **IB15C**, a submicromolar ILT3 binder identified through preliminary structure-activity relationship optimization. Orthogonal validation by microscale thermophoresis, surface plasmon resonance, docking, and site-directed mutagenesis confirmed direct and target-specific ILT3 engagement. Functionally, **IB15C** disrupted the ILT3-ApoE interaction and restored microglial activity in human iPSC-derived microglia, reducing SHP1/2, suppressing cytokine secretion, and enhancing amyloid uptake. **IB15C** also demonstrated favorable in vitro pharmacokinetic and safety properties, supporting further development of ILT3-targeted neuroimmune therapeutics.

---

Establishing direct target engagement of small molecules in a cellular context remains a major challenge in early-stage drug discovery. Although biochemical and biophysical assays enable rapid identification of ligands against purified proteins, these approaches often fail to capture key determinants of intracellular activity, including membrane permeability, protein conformational dynamics, and interactions within native cellular environments.^1-3^ The cellular thermal shift assay (CETSA) provides a label-free approach to address this limitation by quantifying ligand-induced stabilization of proteins in intact cells.^4,5^ Upon thermal challenge, proteins undergo denaturation and aggregation, whereas ligand binding can increase thermal stability, resulting in a measurable shift in the soluble protein fraction.^4-6^ CETSA has been widely applied to confirm target engagement in cells and tissues, and more recently has been adapted to higher-throughput formats suitable for compound profiling and screening.^7-11^

Despite these advances, application of high-throughput CETSA to challenging target classes remains limited. In particular, membrane-associated and non-enzymatic proteins, including immune checkpoint receptors, are difficult to interrogate using conventional biochemical assays due to the absence of catalytic activity and the complexity of their native conformational and interaction states.^12,13^ These challenges highlight the need for scalable, cell-based approaches capable of directly measuring small molecule engagement of such targets in physiologically relevant systems.

Accumulating evidence has established microglia as central drivers of amyloid-β (Aβ) pathology and neuroinflammation in Alzheimer’s disease (AD), with disease progression increasingly linked to signaling pathways that control microglial functional states.^14-18^ Microglial responses are governed by a dynamic balance between activating and inhibitory receptors, including stimulatory pathways such as triggering receptor expressed on myeloid cells 2 (TREM2) and suppressive signaling mediated by CD33 and leukocyte immunoglobulin-like receptor (LILR) family members, all of which regulate phagocytosis, immune activation, and inflammatory tone.^19.20^ Among these inhibitory pathways, leukocyte immunoglobulin-like receptor B4 (LILRB4), also referred to as ILT3, has emerged as an important negative regulator of microglial activity that limits Aβ clearance through ApoE-associated signaling mechanisms.^21-28^

Elevated ILT3 expression has been observed in plaque-associated microglia from AD patient samples and correlates strongly with ApoE abundance as well as pathological hallmarks including phosphorylated tau accumulation.^29^ Mechanistically, ILT3 contains immunoreceptor tyrosine-based inhibitory motifs (ITIMs) that recruit the phosphatases SHP1 and SHP2 upon receptor activation, thereby suppressing cytoskeletal remodeling and phagocytic processes necessary for efficient amyloid clearance.^22,25^ In vivo studies using antibody-based ILT3 blockade have demonstrated enhanced microglial activation, increased Aβ phagocytosis, substantial reductions in amyloid deposition, and attenuation of inflammatory interferon-associated transcriptional programs.^29^ These findings collectively support ILT3 as a compelling neuroimmune checkpoint target in AD and identify the ILT3-ApoE axis as a therapeutically actionable interface. Nevertheless, the development of small molecule modulators for this pathway remains difficult because ILT3 functions through a non-catalytic protein-protein interaction (PPI) surface and because conventional activity-based screening approaches are poorly suited for discovering CNS-penetrant compounds capable of disrupting this signaling network.

Building on our previous work establishing small molecule-based strategies to modulate microglial function through integrated biophysical and cellular screening platforms,^30-33^ we sought to develop a high-throughput and target-engagement-driven discovery pipeline for identifying small molecules targeting ILT3. We employed a high-throughput CETSA (HT-CETSA) approach to directly detect ligand-induced stabilization of ILT3 in a physiologically relevant context. Using this strategy, we identified small molecule modulators of ILT3 that enhance microglial Aβ uptake and suppress inflammatory signaling in human iPSC-derived microglia. Collectively, this work establishes CETSA as an effective platform for discovering small molecule modulators of neuroimmune checkpoints and further supports ILT3 as a therapeutically actionable target in AD.

To establish a target-engagement–based screening platform for identification of ILT3 binders, we developed HT-CETSA using a commercially available CHO-ILT3 stable cell line (Figure 1A). Given that ILT3 functions as a non-enzymatic inhibitory immune receptor lacking intrinsic catalytic activity, CETSA was selected to enable direct monitoring of ligand-induced thermal stabilization under physiologically relevant cellular conditions. The assay workflow consisted of compound incubation in intact cells, thermal challenge, cell lysis, and AlphaLISA-based quantification of soluble ILT3 (Figure 1A).

**Figure 1.**
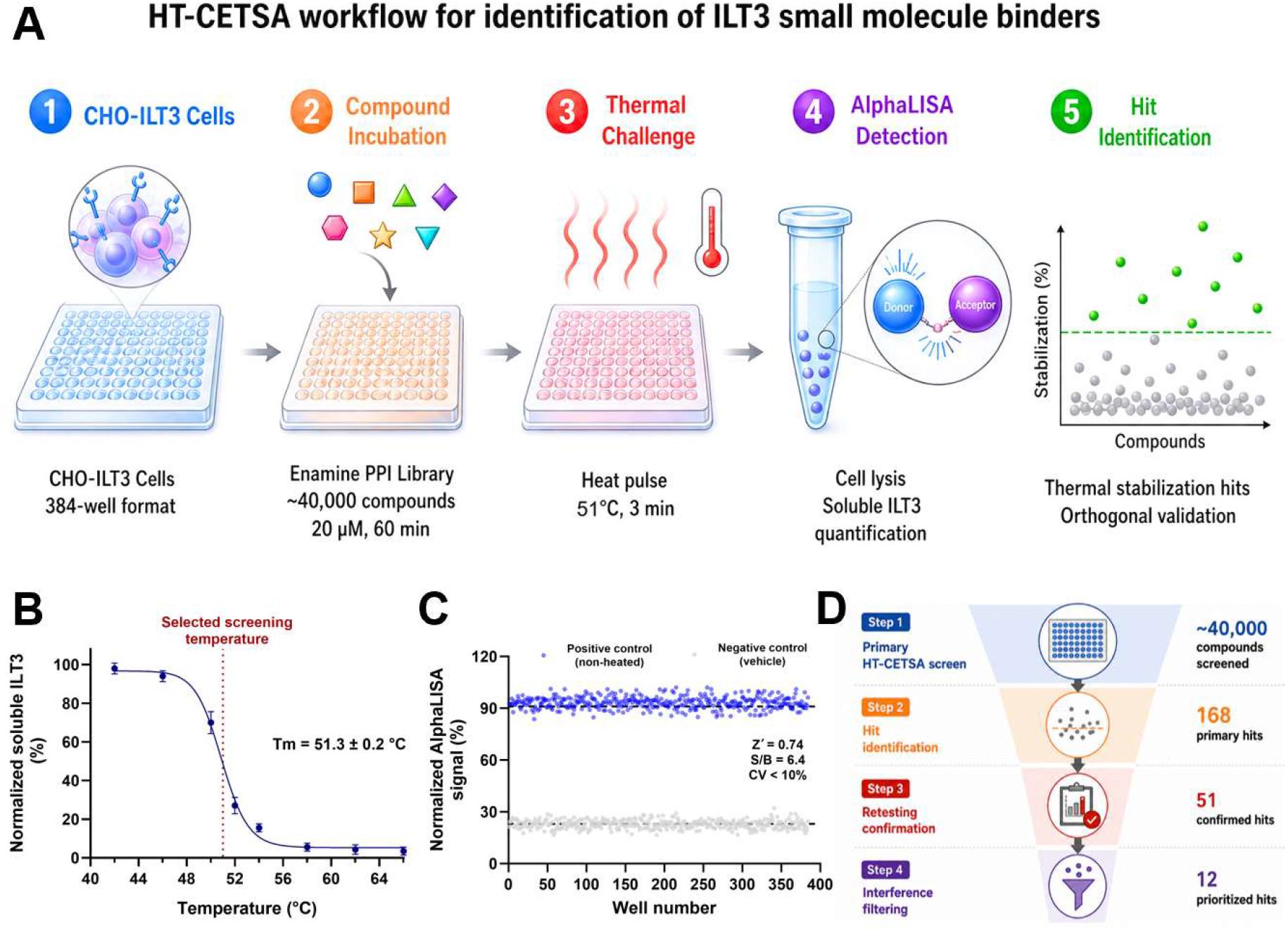
Development and validation of a HT-CETSA platform for identification of ILT3 small molecule binders. **(A)** Schematic overview of the HT-CETSA screening workflow used for identification of ILT3-targeting small molecules. CHO-ILT3 cells were incubated with compounds from the Enamine PPI library followed by thermal challenge, cell lysis, and AlphaLISA-based quantification of soluble ILT3. Compounds inducing thermal stabilization were prioritized for orthogonal validation. **(B)** Thermal denaturation profile of ILT3 in CHO-ILT3 cells determined by HT-CETSA. Cells were exposed to increasing temperatures followed by AlphaLISA quantification of soluble ILT3. The selected screening temperature (51 °C) is indicated by the dashed red line. Data represent mean ± SD (n = 5). **(C)** Assay performance evaluation of the optimized HT-CETSA platform in 384-well format. Positive control (non-heated) and negative control (vehicle-treated, heat-challenged) wells demonstrated robust signal separation. **(D)** Hit identification and triage workflow for prioritization of ILT3 binders. Approximately 40,000 compounds were screened, yielding 168 primary hits, 51 confirmed hits following triplicate retesting, and 12 prioritized hits after removal of PAINS, fluorescent interferers, and aggregation-prone compounds.

We first characterized the thermal denaturation behavior of ILT3 in CHO-ILT3 cells by exposing cells to temperatures ranging from 42-66 °C followed by AlphaLISA quantification of soluble receptor levels after lysis. As shown in Figure 1B, ILT3 exhibited a sharp unfolding transition with an apparent melting temperature (Tm) of 51.3 ± 0.2 °C. Based on the observed denaturation profile, 51 °C was selected as the screening temperature because it provided partial protein unfolding and maximal sensitivity for detecting compound-induced thermal stabilization.

Additional assay optimization was subsequently performed to establish conditions suitable for large-scale screening. A seeding density of 12,000 cells/well in 384-well format yielded optimal assay reproducibility and signal intensity. Compound incubation times between 15 and 120 min were evaluated, with 60 min selected for subsequent studies due to improved stabilization consistency. The assay also demonstrated tolerance to up to 1% DMSO without significant signal deterioration. Under optimized conditions, the HT-CETSA platform demonstrated robust assay performance, yielding an average signal-to-background ratio (S/B) of 6.4 and a mean Z′ factor of 0.74 across validation plates (Figure 1C), supporting suitability for high-throughput screening applications.

Using the optimized platform, approximately 40,000 compounds from the Enamine PPI library were screened at a final concentration of 20 µM. Cells were incubated with compounds for 60 min at 37 °C prior to thermal challenge at 51 °C for 3 min. Following cooling, cell lysis, and AlphaLISA detection, compounds producing stabilization signals greater than 3 standard deviations above the plate median were classified as primary hits. This primary screen identified 168 preliminary hits, corresponding to an overall hit rate of 0.42% (Figure 1D). To evaluate reproducibility, primary hits were retested in triplicate under identical HT-CETSA conditions. Among these, 51 compounds reproducibly induced thermal stabilization of ILT3 and were subsequently advanced for further triaging. Confirmed hits were filtered to remove pan-assay interference compounds (PAINS), fluorescent interferers, and aggregation-prone chemotypes, yielding 12 prioritized hit compounds for downstream validation studies.

To confirm direct target engagement independent of thermal stabilization measurements, the 12 prioritized hits were evaluated using microscale thermophoresis (MST) with recombinant human ILT3 extracellular domain. Eight compounds demonstrated reproducible dose-dependent binding to ILT3, with apparent dissociation constants (Kd) spanning the low-to mid-micromolar range (Table S1). Among these, **IB15** and **IB9** (Figure 2A) emerged as the strongest binders, exhibiting apparent Kd values of 4.9 ± 1.3 µM and 14.2 ± 1.9 µM, respectively (Figure 2B,C). To further confirm direct target engagement using an independent label-free biophysical method, **IB15** was evaluated by surface plasmon resonance (SPR). **IB15** exhibited concentration-dependent binding to recombinant human ILT3 with an apparent Kd of 7.2 ± 0.8 µM (Figure S1), in agreement with the MST-derived affinity and supporting direct interaction with ILT3.

**Figure 2.**
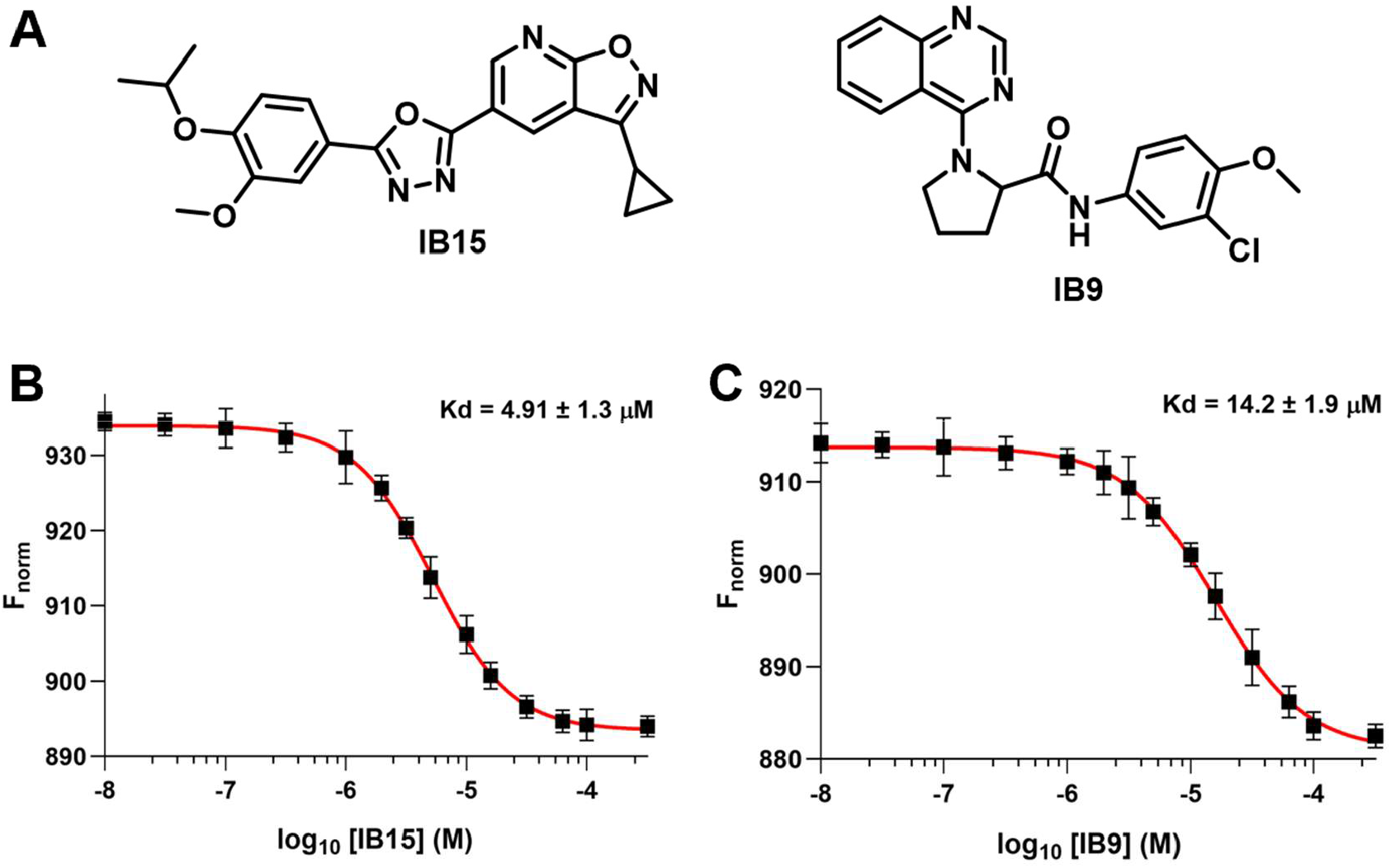
Orthogonal validation of prioritized ILT3 hit compounds by MST. **(A)** Chemical structures of the two top ILT3 hit compounds, **IB15** and **IB9**, identified from the HT-CETSA screening campaign. **(B)** MST binding analysis of **IB15** using recombinant human ILT3 extracellular domain demonstrated dose-dependent target engagement with an apparent dissociation constant (Kd) of 4.9 ± 1.3 µM. **(C)** MST binding analysis of **IB9** confirmed direct interaction with ILT3 with an apparent Kd of 14.2 ± 1.9 µM. Data represent mean ± SD (n=5). Solid red lines represent nonlinear regression fits using a four-parameter variable slope model.

Preliminary catalog-based structure-activity relationship (SAR) analysis revealed that ILT3 binding was highly sensitive to structural modifications around the **IB15** scaffold. Among the evaluated analogs, **IB15** remained one of the most active compounds identified, exhibiting a Kd value of 4.9 ± 1.3 µM. Several derivatives, including **IB15E, IB15F, IB15G**, and **IB15H** showed negligible detectable binding, indicating that replacement of the parent aromatic substituent with more electron-withdrawing, heteroatom-rich, or conformationally altered motifs was poorly tolerated. These findings suggest that preservation of the steric and electronic features of the parent scaffold is important for productive ILT3 engagement.

Interestingly, **IB15J** retained weak but measurable activity despite incorporation of a substantially modified bicyclic substituent, suggesting that limited diversification within this region may still permit partial target interaction. In contrast, highly polar or sulfonamide-containing analogs (e.g., **IB15A**) generally resulted in complete loss of detectable binding, potentially reflecting disruption of favorable hydrophobic interactions or suboptimal positioning within the ILT3 binding interface.

Among the tested analogs, **IB15C** displayed improved affinity relative to the parent scaffold, reaching submicromolar potency (MST Kd = 0.92 ± 0.18 μM), whereas **IB15B** exhibited weaker but measurable engagement. Importantly, the commercially available analog space accessible through the catalog-based SAR approach was limited and did not allow comprehensive or systematic exploration of the scaffold. Consequently, the present SAR analysis should be viewed as preliminary and intended primarily to establish initial structure-activity trends rather than a complete medicinal chemistry optimization campaign. Nevertheless, the observed activity differences across structurally related analogs support a structurally specific mode of ILT3 engagement rather than nonspecific thermal stabilization in the CETSA assay and provide an initial framework for future lead optimization efforts centered around the **IB15** scaffold.

**Table 1.**
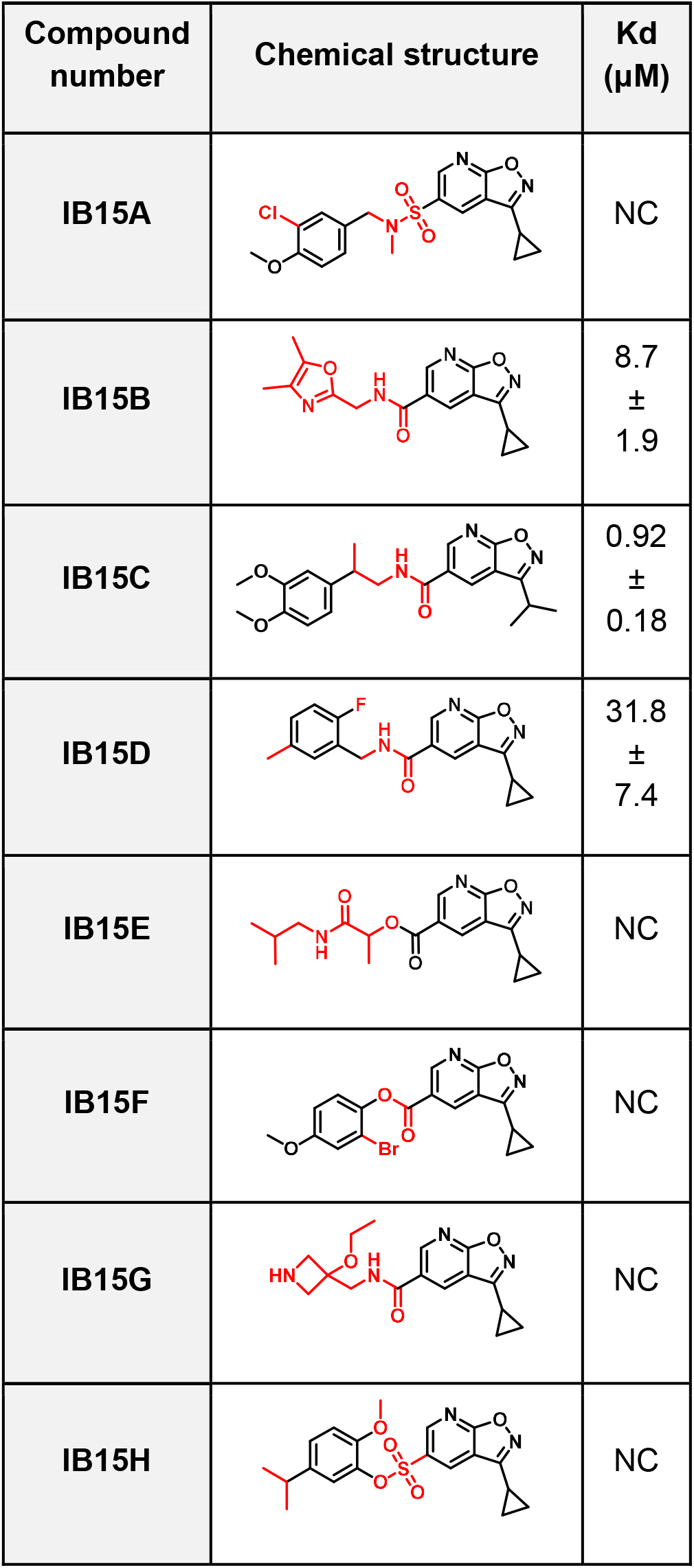
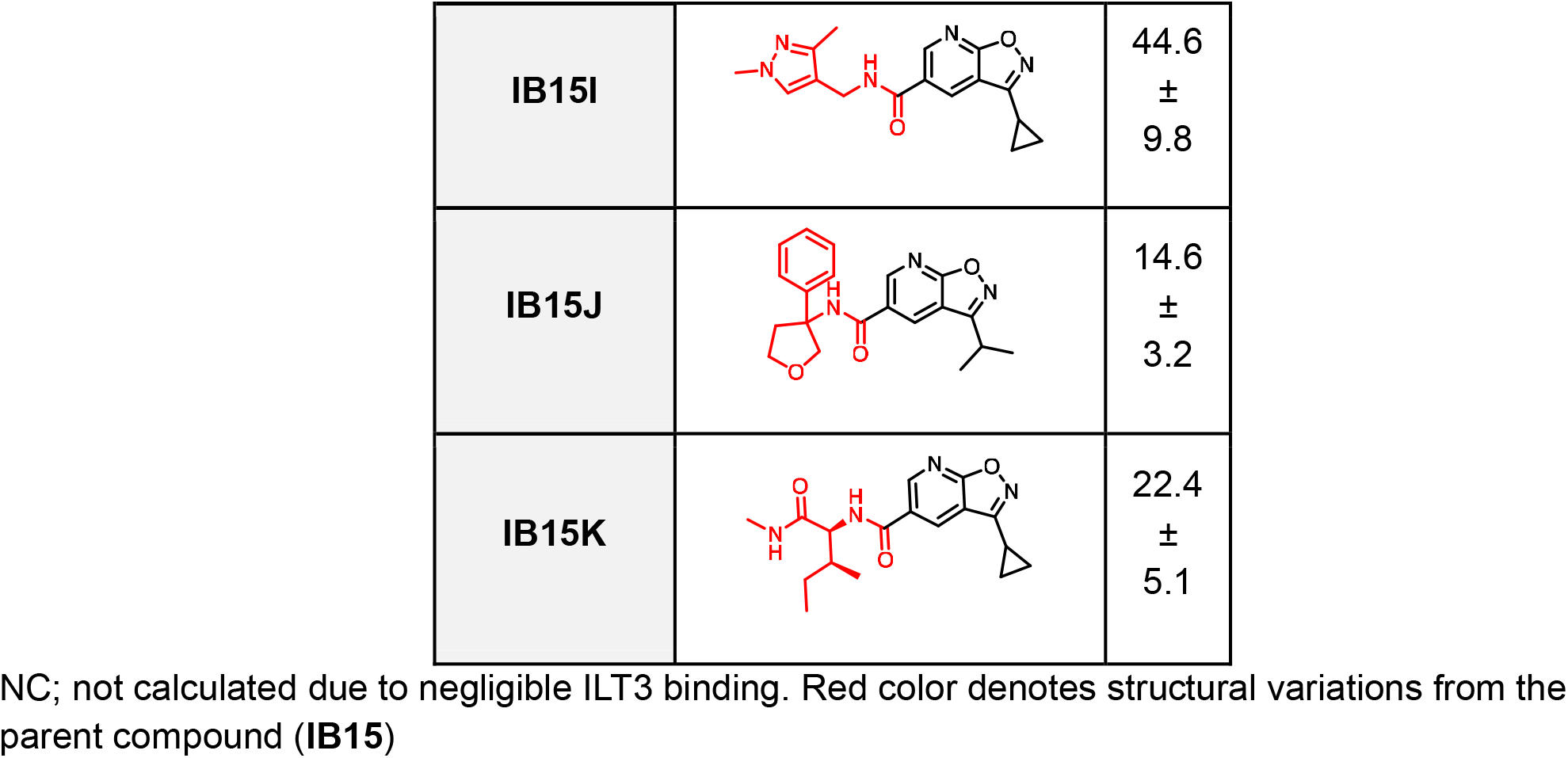
Chemical structures and ILT3 binding affinity (Kd) of IB15 derivatives using MST.

| Compound number | Chemical structure | Kd (μM) |
| --- | --- | --- |
| IB15A |  | NC |
| IB15B |  | 8.7<br>±<br>1.9 |
| IB15C |  | 0.92<br>±<br>0.18 |
| IB15D |  | 31.8<br>±<br>7.4 |
| IB15E |  | NC |
| IB15F |  | NC |
| IB15G |  | NC |
| IB15H |  | NC |
| **IB15I** |  | 44.6<br>±<br>9.8 |
| **IB15J** |  | 14.6<br>±<br>3.2 |
| **IB15K** |  | 22.4<br>±<br>5.1 |

NC; not calculated due to negligible ILT3 binding. Red color denotes structural variations from the parent compound (**IB15**)

To gain structural insight into the binding mode of **IB15C**, the most potent optimized analog identified from catalog-based SAR expansion, molecular docking studies were first performed using the extracellular domain of ILT3. As shown in Figure 3A, 3D docking analysis predicted that **IB15C** occupies a well-defined pocket within ILT3 and adopts a favorable binding orientation stabilized through multiple polar and hydrophobic interactions. The ligand was positioned in close proximity to residues A70, H180, Y181, and L182, which collectively formed a compact interaction network surrounding the compound. Notably, H180 and Y181 appeared centrally involved in ligand stabilization, whereas A70 and L182 contributed more peripheral interactions within the predicted binding pocket.

**Figure 3.**
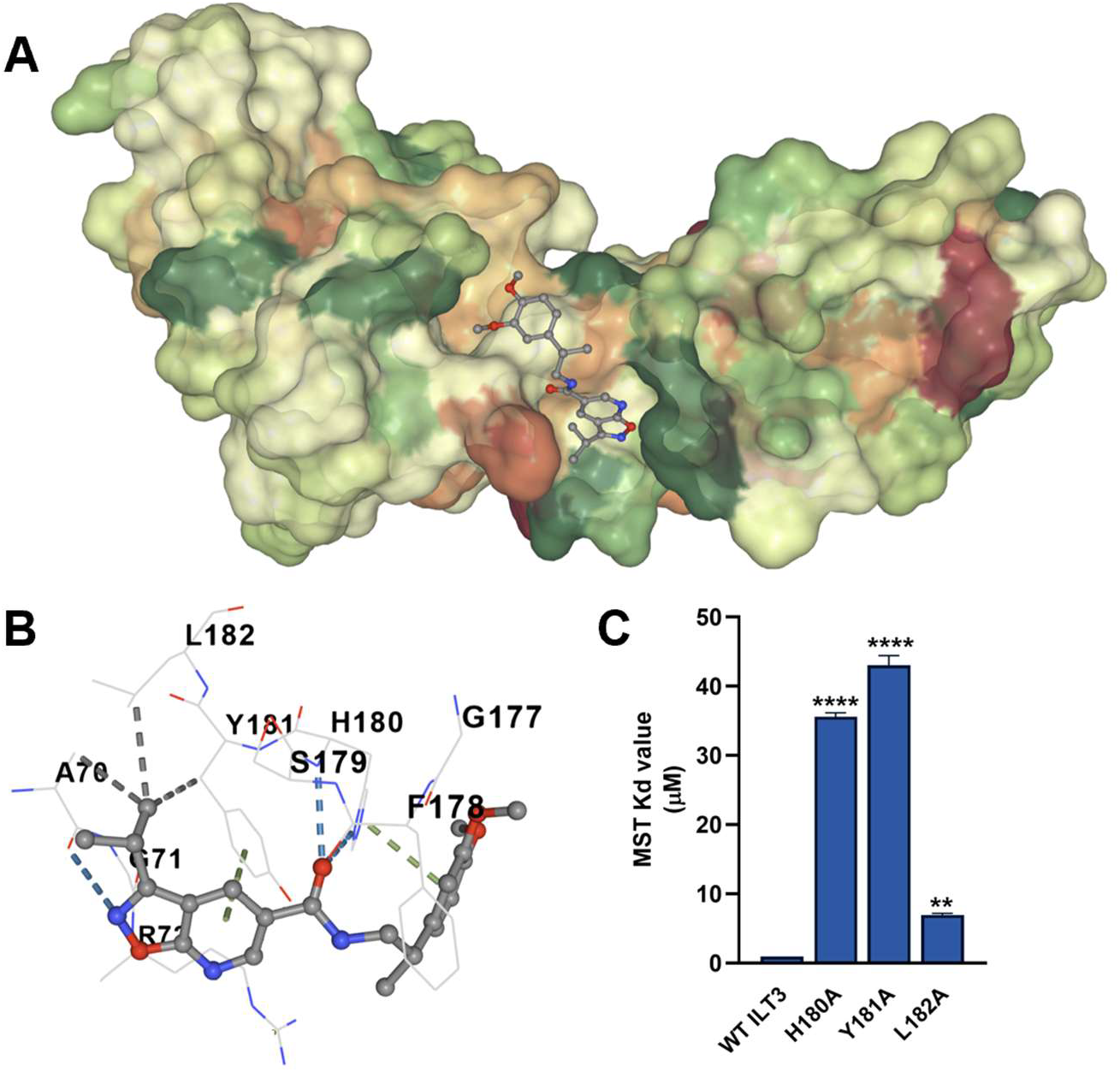
Docking-guided binding mode analysis and site-directed mutagenesis validation of IB15C interaction with ILT3. **(A)** 3D Predicted binding pose of **IB15C** within the ILT3 binding pocket shown on the surface representation of the receptor. **IB15C** occupies a surface-accessible groove enriched in hydrophobic and aromatic residues implicated in ligand recognition. The protein surface is colored based on hydrophobicity. **(B)** Two-dimensional interaction map of the predicted **IB15C**-ILT3 complex. Hydrogen bonding interactions were observed with A70 and H180 (blue dashed lines), while hydrophobic contacts were identified with A70 and L182 (gray dashed lines). In addition, **IB15C** formed π–π stacking interactions with Y181 and H180 (green dashed lines), including two distinct hydrogen bonds with H180. **(C)** Site-directed mutagenesis analysis of residues predicted to contribute to **IB15C** binding. Recombinant ILT3 mutants were evaluated by MST, and Kd values were compared with wild-type (WT) ILT3. Mutations H180A and Y181A substantially impaired **IB15C** binding, whereas L182A produced more moderate effects, supporting the docking-predicted binding mode. Data represent mean ± SD (n=5). Statistical significance was determined relative to WT ILT3.

To further visualize the residue-level interaction network, a 2D interaction map was generated from the docked complex (Figure 3B). This analysis revealed multiple predicted hydrogen bonding and hydrophobic contacts between **IB15C** and ILT3, supporting the proposed binding orientation observed in the 3D model. H180 and Y181 formed prominent interactions with the ligand core, while A70 and L182 contributed additional stabilizing contacts that may help orient the compound within the pocket. Together, these computational analyses suggested that the identified residues could play important roles in mediating **IB15C** binding to ILT3.

To experimentally validate the docking model, site-directed mutagenesis was performed followed by MST binding analysis of alanine-substituted ILT3 variants (Figure 3C). Wild-type ILT3 exhibited high-affinity binding to **IB15C**, whereas mutation of residues within the predicted interaction interface resulted in varying degrees of affinity loss. Among the tested mutants, H180A and Y181A produced the most substantial reductions in binding affinity, with Kd values increasing to ∼35-43 µM relative to wild-type ILT3, indicating that these residues function as critical hotspot interactions for ligand recognition. In contrast, L182A produced more moderate reductions in affinity, with Kd values in the low micromolar range (∼7 µM), suggesting supportive but less dominant contributions to binding. The graded loss of affinity across the mutant panel strongly supports the computationally predicted binding interface and provides orthogonal experimental validation for the proposed **IB15C**-ILT3 interaction model.

Collectively, the combined docking and mutagenesis analyses establish a structurally coherent binding model for **IB15C** and provide mechanistic insight into the molecular determinants governing ILT3 recognition. The pronounced effects associated with H180A and Y181A suggest that these residues form central anchoring interactions required for stable complex formation, whereas L182 appears to contribute secondary stabilizing contacts. These findings further support the conclusion that **IB15C** engages ILT3 through a discrete and target-specific binding pocket, providing a structural foundation for future optimization of ILT3-directed small molecules.

Following confirmation that **IB15C** directly engages ILT3, we next investigated whether ligand binding translated into functional disruption of the ILT3-ApoE interaction. ApoE was selected as the binding partner because of its reported interaction with ILT3 and its established role in regulating neuroimmune signaling in AD. To evaluate the ability of **IB15C** to interfere with receptor-ligand engagement, we employed complementary biochemical assays using both endpoint and real-time binding formats.

In an ELISA-based competition assay, recombinant ILT3 extracellular domain was immobilized and incubated with ApoE in the presence of increasing concentrations of **IB15C**. As shown in Figure 4A, **IB15C** potently inhibited the ILT3-ApoE interaction in a concentration-dependent manner, yielding an IC_50_ of 0.48 ± 0.07 μM. The strong inhibitory activity observed in this assay was consistent with the high-affinity binding profile previously established by MST and supported the ability of **IB15C** to functionally block ligand recognition at the receptor surface.

**Figure 4.**
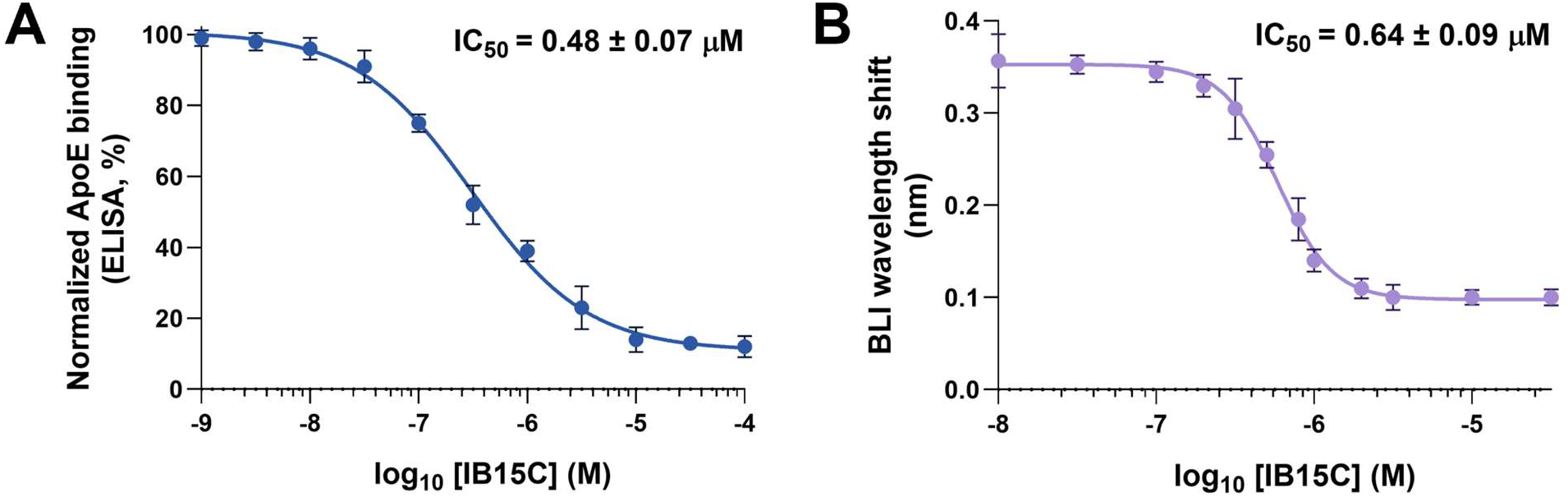
IB15C disrupts the ILT3-ApoE interaction in biochemical and biophysical competition assays. **(A)** ELISA-based competition assay showing concentration-dependent inhibition of ApoE binding to immobilized ILT3 by **IB15C**. Data were fitted using nonlinear regression to determine the IC_50_ value. **(B)** BLI-based competitive binding assay demonstrating concentration-dependent inhibition of the ILT3-ApoE interaction by **IB15C** in real time. Binding responses were analyzed to calculate IC_50_ values. Data are presented as mean ± SEM (n=5).

To further validate these findings using an orthogonal biophysical approach, competitive binding experiments were performed by biolayer interferometry (BLI), enabling real-time monitoring of ApoE association with ILT3. Consistent with the ELISA results, **IB15C** produced a robust concentration-dependent reduction in binding signal, with an IC_50_ of 0.64 ± 0.09 nM (Figure 4B). The close agreement between ELISA and BLI measurements across independent assay platforms strongly supports a direct mechanism in which **IB15C** disrupts ILT3-ApoE complex formation rather than indirectly altering assay components or protein stability.

Together, these findings demonstrate that **IB15C** not only binds ILT3 with high affinity but also effectively blocks interaction with its endogenous ligand ApoE. The concordance between orthogonal assay systems further strengthens the evidence for target-specific functional inhibition and supports the utility of **IB15C** as a small molecule modulator of the ILT3 signaling axis.

To determine whether biochemical inhibition of the ILT3-ApoE interaction translates into functional rescue in disease-relevant cellular systems, **IB15C** was evaluated in human iPSC-derived microglia exposed to Aβ-driven inflammatory conditions. Microglial cultures were stimulated with Aβ_42_ oligomers in the presence or absence of ApoE to induce ligand-dependent ILT3 signaling, followed by treatment with **IB15C** at 1, 5, or 10 µM.

We first investigated early signaling events downstream of ILT3 activation by assessing phosphorylation of the inhibitory phosphatases SHP1 and SHP2, which are recruited to ITIM-containing receptors following ligand engagement. Exposure to Aβ_42_ together with ApoE markedly increased phospho-SHP1 and phospho-SHP2 levels relative to vehicle-treated cells and cultures stimulated with Aβ_42_ alone (Figures 5A and 5B), consistent with activation of ILT3-associated inhibitory signaling pathways. Treatment with **IB15C** reduced SHP1 and SHP2 phosphorylation in a concentration-dependent manner, with partial suppression observed at 1 µM and substantially greater inhibition at 5 and 10 µM (Figures 5A and 5B). These findings indicate that **IB15C** effectively interferes with ligand-driven ILT3 signaling in human microglia.

**Figure 5.**
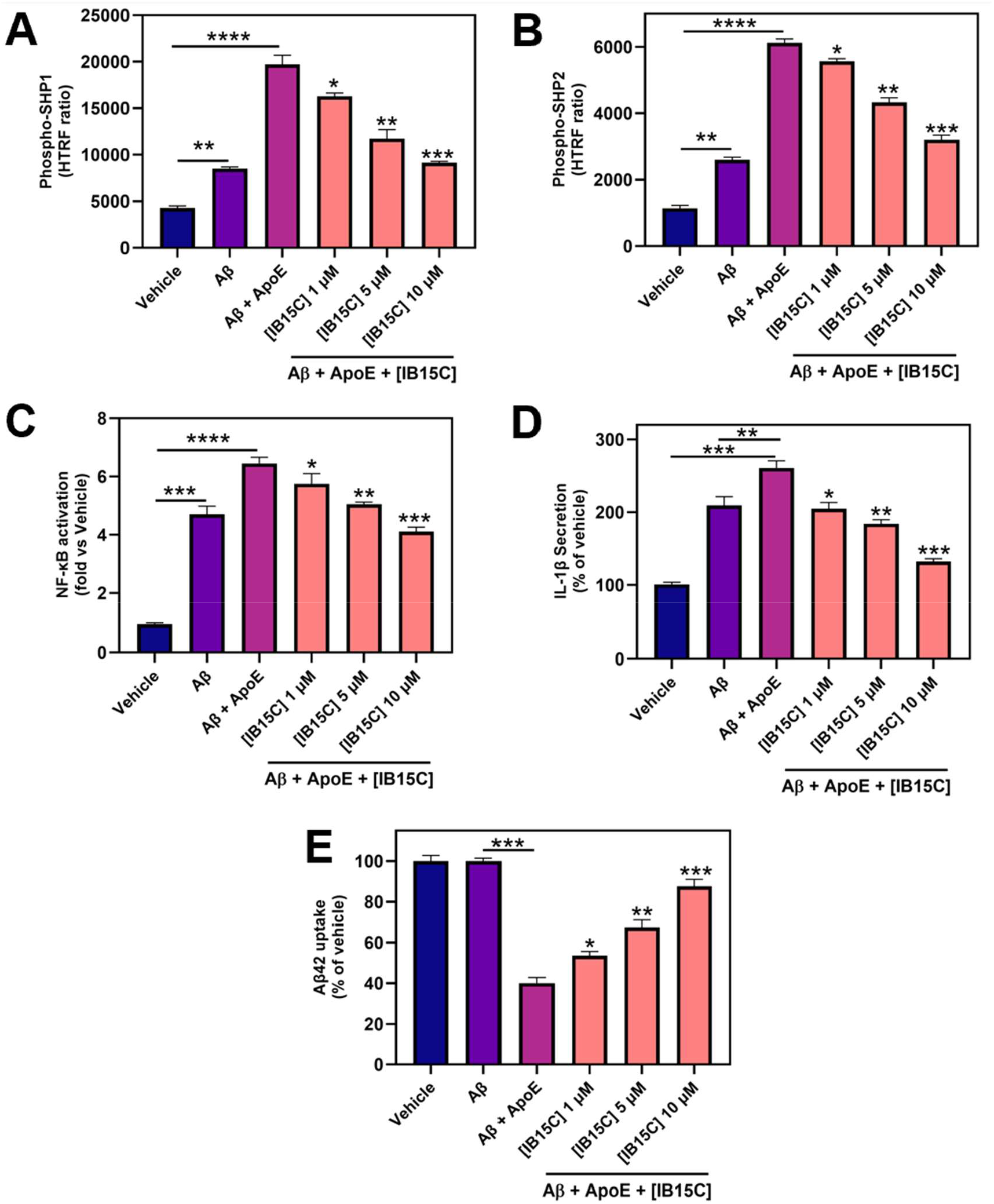
IB15C suppresses ILT3-dependent signaling, reduces inflammatory activation, and restores Aβ uptake in human iPSC-derived microglia. **(A**,**B)** Phosphorylation of SHP1 **(A)** and SHP2 **(B)** following stimulation with Aβ_42_ and ApoE. ApoE enhanced SHP1/2 phosphorylation relative to Aβ_42_ alone, consistent with ILT3 pathway activation, while **IB15C** reduced phospho-SHP1 and phospho-SHP2 levels in a dose-dependent manner. **(C)** NF-κB activation relative to vehicle control. ApoE further potentiated Aβ_42_-induced NF-κB signaling, which was attenuated by **IB15C. (D)** IL-1β secretion normalized to vehicle. **IB15C** decreased ApoE-enhanced inflammatory cytokine release in a concentration-dependent manner. **(E)** Aβ_42_ uptake by microglia. ApoE impaired amyloid uptake, whereas **IB15C** restored uptake in a dose-dependent fashion. Data are presented as mean ± SD (n = 5). Statistical significance was determined by one-way ANOVA with multiple-comparisons testing. \*\**p* < 0.01, \*\*\**p* < 0.001, \*\*\*\**p* < 0.0001; *ns*, not significant.

We next examined downstream inflammatory signaling by monitoring NF-κB activation. Stimulation with Aβ_42_ increased NF-κB activity, and this response was further potentiated by ApoE co-treatment (Figure 5C). Administration of **IB15C** attenuated NF-κB activation in a dose-responsive manner, with the strongest suppression observed at 10 µM, where signaling approached near-basal levels (Figure 5C). The coordinated reduction in SHP1/2 phosphorylation and NF-κB activity supports a mechanistic relationship between ILT3 engagement and downstream inflammatory signaling cascades. To evaluate functional inflammatory responses, secretion of IL-1β was quantified in conditioned media. Aβ_42_ stimulation elevated IL-1β release, and the addition of ApoE further amplified cytokine production (Figure 5D). **IB15C** significantly reduced IL-1β secretion across all tested concentrations, with more pronounced effects at 5 and 10 µM, consistent with inhibition of upstream ILT3 signaling events. We next assessed microglial phagocytic capacity by measuring Aβ uptake. Whereas Aβ_42_ alone produced minimal effects on internalization, co-treatment with ApoE substantially impaired Aβ uptake (Figure 5E), reflecting suppression of microglial clearance activity. **IB15C** restored Aβ internalization in a concentration-dependent fashion, with significant recovery observed at 5 and 10 µM (Figure 5E).

Importantly, **IB15C** did not produce detectable cytotoxicity under any treatment condition, as cell viability remained unchanged across all tested concentrations (Figure S2). Collectively, these results demonstrate that pharmacological inhibition of ILT3-ApoE signaling by **IB15C** reverses inhibitory microglial signaling and restores functional responses under Aβ-associated inflammatory conditions.

To further evaluate the translational potential of **IB15C**, a comprehensive in vitro pharmacokinetic (PK) and developability assessment was performed (Table 2). **IB15C** exhibited a balanced physicochemical profile with a LogD7.4 value of 2.9, consistent with moderate lipophilicity and favorable drug-like properties. The compound demonstrated high aqueous solubility, with kinetic solubility reaching 95 µM in 1% DMSO/PBS and FaSSIF solubility of 128 µM, indicating good compatibility with physiologically relevant intestinal conditions and supporting oral developability.

Assessment of membrane permeability revealed that **IB15C** possesses favorable passive diffusion properties. In the PAMPA-BBB assay, the compound displayed a permeability coefficient of 4.2 × 10^-6^ cm/s, suggesting the capacity for blood–brain barrier penetration. Consistent with these findings, MDCK-MDR1 transport studies demonstrated measurable bidirectional permeability with an efflux ratio of 1.51, indicating only modest susceptibility to active efflux mechanisms. Together, these data support adequate cellular permeability while suggesting that **IB15C** is not a strong substrate for MDR1-mediated transport.

**IB15C** also demonstrated acceptable metabolic stability across multiple assay systems. In liver microsomes, the compound exhibited moderate stability in both mouse and human preparations, with half-lives of 29.8 ± 2.1 min and 46.5 ± 3.4 min, respectively. Corresponding intrinsic clearance values further supported a metabolically manageable profile. In addition, **IB15C** showed strong stability in plasma and buffer systems, with prolonged half-lives in mouse plasma, human plasma, and PBS, indicating favorable chemical and biological stability under physiologically relevant conditions. The compound also remained highly stable in simulated gastric and intestinal fluids, with >89% remaining after 2 h incubation, supporting its compatibility with oral exposure conditions.

Importantly, **IB15C** exhibited a favorable preliminary safety profile. No measurable cytotoxicity was observed in WI-38 or HS27 human cell lines at concentrations up to 100 µM. Likewise, the compound showed minimal hERG channel inhibition, with an IC_50_ greater than 100 µM, suggesting low risk for cardiac liability. CYP inhibition profiling further demonstrated limited inhibition across major drug-metabolizing isoforms, including CYP3A4, CYP2D6, CYP2C9, and CYP2C19, with all values remaining below 10% inhibition at 10 µM. Collectively, these findings indicate that **IB15C** combines functional ILT3 inhibition with a favorable in vitro PK and safety profile, supporting its suitability as a promising small molecule scaffold for further optimization and in vivo evaluation.

**Table 2.** In vitro PK profile of **IB15C**.

| PK parameter | Value | PK parameter | Value |
| --- | --- | --- | --- |
| LogD<sub>7.4</sub><sup>a</sup> | 2.9 | t<sub>1/2</sub> [min] mouse plasma | 112 |
| Kinetic Solubility 1%<br>DMSO/PBS [μM]<sup>b</sup> | 95 | t<sub>1/2</sub> [min] human plasma | 156 |
| Solubility in FaSSIF [μM] | 128 | t<sub>1/2</sub> [min] PBS | 241 |
| PAMPA-BBB P<sub>e</sub> [cm/s] | $4.2 \times 10^{-6}$ | PPB human [%]<sup>c</sup> | $96.8 \pm 0.53$ |
| MDCK–MDR1 P<sub>app</sub><br>(A→B) | $1.15 \times 10^{-5}$ | IC<sub>50</sub> against WI-38<br>[μM] | >100 |
| MDCK–MDR1 $P_{app}$<br>(B→A) | $1.74 \times 10^{-5}$ | IC<sub>50</sub> against HS 27<br>[μM] | >100 |
| MDCK–MDR1 Efflux<br>Ratio (B→A/A→B) | 1.51 | IC<sub>50</sub> against hERG<br>[μM] | >100 |
| Stability in simulated<br>gastric fluid (pH 1.2, 2h) | 91% remaining | CYP3A4 inhibition (%)<br>at 10 μM) | 7.4% |
| Stability in simulated<br>intestinal fluid (pH 6.8,<br>2h) | 89% remaining | CYP2D6 inhibition (%)<br>at 10 μM) | 6.1% |
| $t_{1/2}$ [min] mouse liver<br>microsomes<sup>c</sup> / CL<sub>int</sub><sup>d</sup> | $29.8 \pm 2.1 / 45.6$<br>$\pm 3.3$ | CYP2C9 inhibition (%)<br>at 10 μM) | 3.4% |
| $t_{1/2}$ [min] human liver<br>microsomes<sup>c</sup> / CL<sub>int</sub><sup>d</sup> | $46.5 \pm 3.4 / 29.2$<br>$\pm 2.4$ | CYP2C19 inhibition (%)<br>at 10 μM) | 2.7% |

<sup>a</sup>LogD (pH = 7.4) the distribution coefficient at physiological pH 7.4. <sup>b</sup>Kinetic solubility measured in 1% DMSO/PBS (pH 7.4) by UV–vis spectrophotometry (n=3). <sup>c</sup>Values are mean ± SD (n=3). <sup>d</sup>Intrinsic clearance expressed in $\mu\text{L}/(\text{min} \times \text{mg})$ protein. Values are mean ± SD (n=3).

In conclusion, this study establishes a high-throughput CETSA-based discovery platform for identification of small molecule modulators targeting the inhibitory immune receptor ILT3 (LILRB4). Using this target-engagement-driven approach, we identified and optimized **IB15C**, a submicromolar ILT3 binder that demonstrated direct target engagement across multiple orthogonal biophysical assays, including MST and SPR. Docking-guided mutagenesis studies further supported a structurally defined and target-specific binding mode. Importantly, **IB15C** functionally disrupted the ILT3-ApoE interaction and restored microglial activity in human iPSC-derived microglia under Aβ-associated inflammatory conditions, leading to reduced inhibitory signaling, suppression of pro-inflammatory cytokine production, and enhanced Aβ uptake. In addition, **IB15C** exhibited favorable in vitro PK and preliminary safety properties consistent with further therapeutic development. Collectively, these findings establish ILT3 as a tractable neuroimmune checkpoint for small molecule intervention in AD and highlight HT-CETSA as a scalable strategy for discovering ligands against challenging non-enzymatic immune receptors.

## Notes

### Competing Interest Statement

The authors have declared no competing interest.

