## Supporting Information for "High-Throughput CETSA Identifies Small Molecule Modulators of ILT3 (LILRB4) with Functional Activity in Human iPSC-Derived Microglia for Alzheimers Disease"

*Electronic Supplementary Information for*

AUTHOR ADDRESS

* To whom correspondence should be addressed:

| **Contents** |  |
| --- | --- |
| Experimental  Binding affinities of validated hits as LILRB4 binders measured by MST | S3  S7 |
| SPR binding analysis of **IB15** against recombinant human ILT3 | S8 |
| Cell viability analysis in human iPSC-derived microglia following treatment with **IB15C** under the indicated experimental conditions | S8 |

**Experimental:**

**High-Throughput Cellular Thermal Shift Assay (HT-CETSA).** CHO cells stably expressing human ILT3/LILRB4 (from Creative Biogene, Cat# CSC-RO0667) were maintained in Ham’s F-12 medium supplemented with 10% fetal bovine serum and 1% penicillin-streptomycin under standard humidified culture conditions (37 °C, 5% CO_2_). For HT-CETSA experiments, cells were seeded into white 384-well assay plates at a density of 12,000 cells/well in a final assay volume of 40 µL and allowed to adhere overnight.

Compounds from the Enamine PPI library (40,640 compounds) were acoustically dispensed into assay plates using an Echo acoustic liquid handler to achieve a final concentration of 20 µM with 1% (v/v) DMSO. Following compound addition, cells were incubated for 60 min at 37 °C to permit intracellular target engagement. Non-heated wells were included as positive controls representing maximal soluble ILT3 signal, whereas heat-challenged DMSO-treated wells served as negative controls.

For thermal denaturation profiling, assay plates were sealed and heated over a temperature range of 42-66 °C for 3 min using a calibrated Bio-Rad C1000 Touch thermal cycler, followed by equilibration at room temperature for 3 min. For screening experiments, an isothermal format was employed using a fixed challenge temperature of 51 °C, selected based on the experimentally determined ILT3 melting profile. Following thermal challenge, cells were lysed directly in assay plates by addition of lysis buffer supplemented with protease inhibitor cocktail and incubated for 30 min at 4 °C with gentle agitation. Lysates were subsequently clarified by centrifugation at 4000 × g for 10 min to remove aggregated and insoluble protein species generated during thermal denaturation.

The soluble fraction of ILT3 remaining after thermal challenge was quantified using an AlphaLISA-based immunodetection format. Briefly, clarified lysates were incubated with ILT3-specific biotinylated detection antibody and AlphaLISA acceptor beads according to the manufacturer’s instructions, followed by addition of streptavidin-coated donor beads under reduced-light conditions. After incubation at room temperature, AlphaLISA signal was measured using Tecan Spark plate reader. For assay validation, signal-to-background ratio (S/B) and Z′ factor were calculated from positive and negative control wells across validation plates. Under optimized conditions, the assay yielded an average S/B ratio of 6.4 and mean Z′ factor of 0.74, supporting suitability for high-throughput screening applications.

For primary screening, compounds from the Enamine PPI library were screened at a final concentration of 20 µM in 384-well format under isothermal HT-CETSA conditions using a fixed thermal challenge temperature of 51 °C. Each assay plate contained non-heated control wells representing maximal soluble ILT3 signal and heat-challenged DMSO-treated wells representing baseline denatured protein signal.

Following AlphaLISA detection, raw luminescence values were normalized using the following equation:

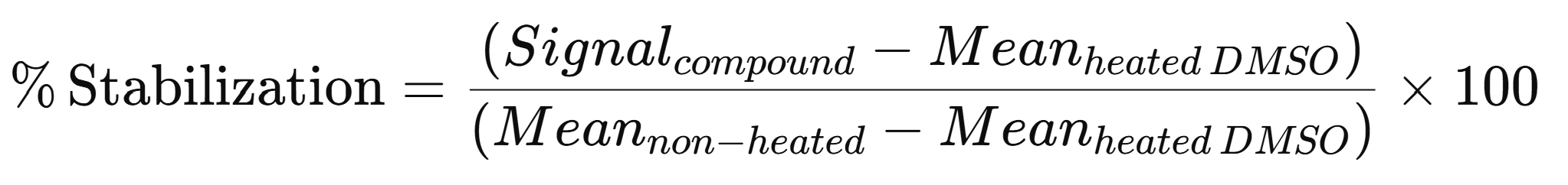

where Signal_compound_ represents the AlphaLISA signal obtained from compound-treated heated wells, Mean_heated DMSO_ corresponds to the average signal from heat-challenged vehicle control wells, and Mean_non−heated_ represents the average signal from non-heated control wells.

Compounds producing stabilization values greater than three standard deviations above the plate median were classified as primary hits. In parallel, assay robustness was continuously monitored using signal-to-background ratio (S/B) and Z′ factor calculations, with plates displaying Z′ values below 0.5 excluded from analysis.

Primary hit compounds were cherry-picked and retested in triplicate under identical HT-CETSA conditions. Compounds demonstrating reproducible thermal stabilization across replicate experiments were advanced for secondary triaging. Structural and assay-behavior filters were subsequently applied to remove pan-assay interference compounds (PAINS), fluorescent interferers, and aggregation-prone chemotypes. Remaining prioritized hits were then subjected to MST biophysical validation.

**MST analysis for ILT3 binding.** Binding affinities of selected hits were quantified by microscale thermophoresis (MST) using a Monolith X instrument (NanoTemper Technologies). His-tagged ILT3 protein (30 nM) was labeled with RED-tris-NTA 2nd Generation dye using the Monolith His-Tag Labeling Kit (Cat. #MO-L018) following the manufacturer’s guidelines. Compound titrations were prepared as serial dilutions (PBS buffer, pH 7.4, 0.05% Tween-20, 1% (v/v) DMSO). Following a 15 min incubation at room temperature in the dark, samples were loaded into Monolith capillaries (Cat. #MO-K022) and analyzed at 25 °C using 40% LED power and medium MST power settings. Normalized fluorescence (F_norm_) values were determined as the ratio of fluorescence intensity after and before IR laser heating. Each compound was evaluated in five technical replicates. Dissociation constants (Kd) were calculated from three independent experiments using GraphPad Prism 10. Kd values are reported as mean ± SD (n=5).

**SPR analysis.** SPR validation of ILT3 binding was performed using Biacore™ 8K SPR system (Cytiva, Marlborough, MA, USA). ILT3-His Protein (10 μg/mL in PBS, pH 7.4) was immobilized on a Series S Sensor Chip CM5 (29104988, Cytiva, Marlborough, MA, USA) using a commercial amine coupling kit (BR100050, Cytiva, Marlborough, MA, USA). Immobilization was performed at a flow rate of 10 μL/min for 420 s, followed by blocking with ethanolamine. The flow cell with solely ethanolamine-block on the same channel served as the reference. Immobilization buffer: PBS-P+ (28995084, Cytiva, Marlborough, MA, USA). Briefly, gradient concentrations of the compound were prepared in the assay buffer, and injected over the sensor chip in a single-cycle kinetics model with a flow rate at 30 μL/min for 120 s per injection. After each injection, a 30 s-regeneration was performed at a flow rate of 30 μL/min using the regeneration buffer. Assays were performed in triplicate, and Kd values are reported as mean ± SD (n=3).

**Computational studies**. Molecular docking studies were performed to characterize the interactions of **IB15C** with ILT3. The crystal structure of ILT3 (PDB ID: 3P2T) was obtained from the Protein Data Bank. The solvents and water were removed from the receptor prior to docking. The lead compounds in SMILES format were converted into .mol2 and subsequently to .pdbqt using openbabel and MGLTools. The pdbqt file includes Gasteiger charges and information on rotatable bonds. We have carried out molecular docking using AutoDock4.0.

**ELISA-based competition assay.** ILT3 extracellular domain (2 µg/mL) was immobilized on 96-well high-binding plates (Corning) overnight at 4°C. Plates were blocked with 5% BSA in PBS for 1 h at room temperature. Recombinant human ApoE (R&D Systems) was added at a fixed concentration (100 nM) in the presence of serial dilutions of compounds. Bound ApoE was detected using an HRP-conjugated anti-ApoE antibody, followed by TMB substrate development. Absorbance was measured at 450 nm using a Tecan plate reader. Data were normalized to DMSO controls and fitted to determine IC_50_ values using nonlinear regression (GraphPad Prism v10). Reported values represent mean ± SD (n=5).

**BLI competition study.** BLI experiments were performed using an Octet RED96 system (Sartorius). LILRB4 protein was immobilized onto Ni-NTA biosensors, followed by equilibration in assay buffer (PBS + 0.05% Tween-20). ApoE (100 nM) was associated in the presence of increasing concentrations of compounds. Binding responses were monitored as wavelength shifts, and inhibition curves were generated to determine IC_50_ values. Data were analyzed using Octet Data Analysis software. Reported values represent mean ± SD (n=5).

**Site-directed mutagenesis.** Alanine substitutions were introduced into the human ILT3 extracellular domain using the QuikChange Lightning Site-Directed Mutagenesis Kit (Agilent Technologies) according to the manufacturer’s protocol. Briefly, mutagenic primers were designed to incorporate the desired codon substitutions and PCR amplification was performed using a high-fidelity polymerase. Parental methylated DNA was digested with DpnI, and the resulting plasmids were transformed into *E. coli* DH5α cells. All mutations were confirmed by Sanger sequencing. Recombinant wild-type and mutant ILT3 proteins were expressed and purified as described above. Protein integrity and purity (>90%) were verified by SDS-PAGE and analytical size-exclusion chromatography. Binding affinities of mutant proteins were evaluated by MST under identical conditions to wild-type controls, and Kd values were obtained from nonlinear regression fits as described above.

**Human iPSC-derived microglia assays.** Human iPSC-derived microglia (FujiFilm Cellular Dynamics) were cultured in microglia maintenance medium (Axol Bioscience Ltd, Catalog #ax0660) supplemented according to the supplier’s recommendations at 37 °C in a humidified 5% CO_2_ incubator. Cells were plated at 50,000 cells per well in poly-D-lysine-coated 96-well plates and allowed to recover for 24 h prior to treatment. Aβ42 oligomers were prepared by dissolving lyophilized Aβ42 peptide (≥95% purity; AnaSpec, Catalog# AS-20276) in HFIP, followed by evaporation, resuspension in DMSO, and oligomerization in PBS at 4 °C for 24 h. Cells were treated with Aβ42 (1 μM) in the presence or absence of recombinant human ApoE (100 nM), followed by addition of **IB15C** at indicated concentrations (1-10 μM). Treatments were performed for 24 h. Vehicle controls contained equivalent DMSO concentrations (<0.5%).

**Signaling assays.** Phosphorylation of SHP1 and SHP2 was quantified using phospho-specific ELISA kits (abcam, catalog# ab279924 and ab314344, respectively) following the manufacturer’s protocol. Briefly, cells were lysed in ice-cold lysis buffer supplemented with protease and phosphatase inhibitors. Lysates were normalized for total protein content (BCA assay), and equal amounts were loaded per well. Reported values represent mean ± SD (n=5).

NF-κB activation was measured using a luciferase reporter assay in human iPSC-derived microglia. Cells were plated at 50,000 cells per well in white 96-well plates and transfected with an NF-κB firefly luciferase reporter (Promega) and a Renilla luciferase control plasmid (Promega) using Lipofectamine 3000 (Thermo Fisher Scientific, Cat. #L3000008). Twenty-four hours post-transfection, cells were treated with Aβ42 oligomers (1 μM) ± ApoE (100 nM) and **IB15C** (1-10 μM) for 24 h. Luciferase activity was measured using the Dual-Glo Luciferase Assay System (Promega, Cat. #E2920). NF-κB activity was calculated as the ratio of firefly to Renilla signal and expressed as fold change relative to vehicle controls. Reported values represent mean ± SD (n=5).

**Cytokine measurements.** IL-1β levels in culture supernatants were quantified using a human IL-1β ELISA kit (R&D Systems, catalog# DLB50). Supernatants were collected, clarified by centrifugation (1,000 × g, 5 min), and analyzed according to the manufacturer’s instructions. Cytokine concentrations were interpolated from standard curves and normalized where appropriate. Reported values represent mean ± SD (n=5).

**Aβ uptake assay.** Fluorescently labeled Aβ42 (HiLyte Fluor 555-conjugated from AnaSpec, catalog# AS-60480-01) was added to cells at a final concentration of 500 nM following treatment conditions described above. After incubation (4 h), cells were washed extensively with PBS and treated with trypan blue to quench extracellular fluorescence. Intracellular fluorescence was quantified using a Tecan Spark plate reader and normalized to cell number (total protein).

**Cell viability.** Cell viability was assessed using the CellTiter-Glo luminescent assay (Promega, catalog#G7570) according to the manufacturer’s protocol. Following treatment, reagent was added directly to wells, incubated for 10 min at room temperature, and luminescence was measured. Viability was normalized to vehicle-treated controls. Reported values represent mean ± SD (n=5).

**PK studies.** In vitro PK profiling was performed as we previously reported.^1^

**Table 1. Binding affinities of validated hits as ILT3 binders measured by MST.**
Equilibrium dissociation constants (Kd, µM) for the compounds were determined using microscale thermophoresis (MST).

| **Comp. ID** | **Enamine Code** | **SMILES** | **MST KD (μM)** |
| --- | --- | --- | --- |
| **IB15** | Z1170449523 | COC=1C=C(C=CC1OC(C)C)C2=NN=C(O2)C3=CN=C4ON=C(C4=C3)C5CC5 | 4.9 ± 1.3 |
| **IB9** | Z223591924 | COC=1C=CC(=CC1Cl)NC(=O)C2CCCN2C=3N=CN=C4C=CC=CC34 | 14.2 ± 1.9 |
| **IB83** | Z235353857 | CCC=1C=CC(=CC1)N2C(=O)C3N=NN(CC(=O)NC=4C=CC=C(F)C4)C3C2=O | 25.7 ± 2.8 |
| **IB161** | Z1170429373 | FC(F)(F)C=1C=CN=C(N1)N2CCC3(CC3C4=NN=C5CCCCCN45)CC2 | 27.5 ± 3.6 |
| **IB127** | Z1318076219 | CC1=CC(CN2CC(C)CN(CC=3C=C(C)ON3)C4=CC=CC=C42)=NO1 | 51.4 ± 3.4 |
| **IB48** | Z1230705474 | CC=1C=C(NC(=O)C2C(=O)N(C)N=C2C)N(CC=3C=CC=CC3)N1 | 58.7 ± 4.1 |
| **IB59** | Z141018980 | O=C(CCC1=CNC=2C=CC=CC12)N3CCCC3C=4C=CC=5OCCCOC5C4 | 66.5 ± 4.8 |
| **IB31** | Z1166112132 | FC=1C=CC=CC1C2=NN=C(O2)C3(CCOCC3)C=4C=CC=5OCOC5C4 | 93.7 ± 5.3 |

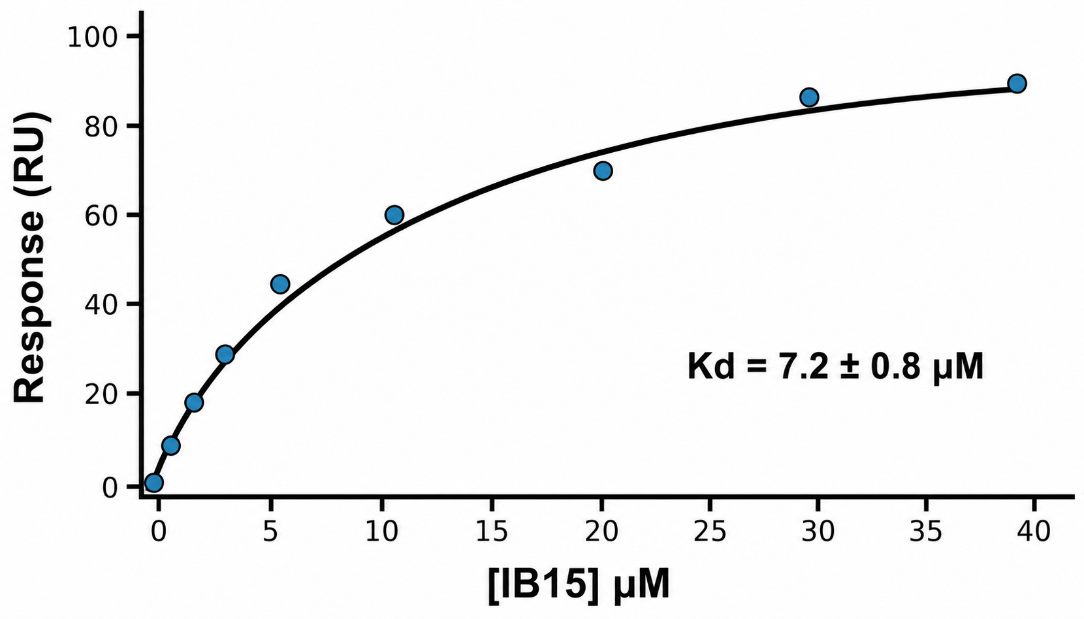

**Figure S1.** **SPR binding analysis of IB15 against recombinant human ILT3.** Increasing concentrations of **IB15** produced a concentration-dependent binding response, consistent with direct interaction with ILT3. Nonlinear fitting of the equilibrium binding response yielded an apparent dissociation constant (Kd) of 7.2 ± 0.8 µM (n=3). Data points represent experimental measurements, and the solid black line indicates the fitted binding isotherm.

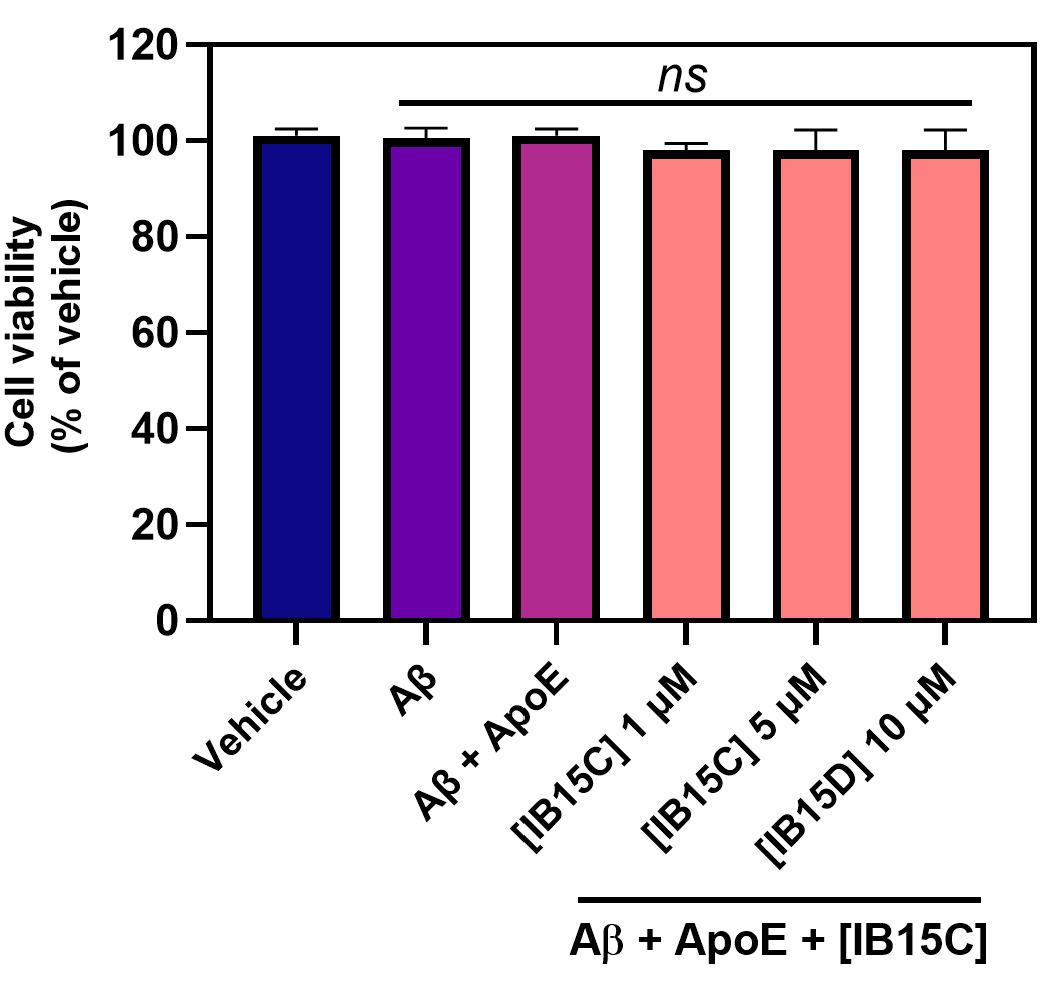

**Figure S2. Cell viability analysis in human iPSC-derived microglia following treatment with IB15C under the indicated experimental conditions.** No significant changes in viability were observed across all groups, indicating that the biological effects of **IB15C** were not associated with cytotoxicity. Data are presented as mean ± SD (n = 5). Statistical analysis was performed using one-way ANOVA with multiple-comparisons testing; *ns*, not significant.
